# A compartmentalised cochlear ageing programme: weak systemic concordance and a distinct macrophage compartment

**DOI:** 10.64898/2026.11.17.767463

**Authors:** Zhengyi Tang, Ting Liao, Qiuping Lin

## Abstract

Whether the ageing cochlea follows the systemic ageing programmes defined by multi-organ atlases is unknown. We integrated single-nucleus RNA-sequencing data for 13,988 mouse cochlear nuclei (3, 12 and 24 months) with the Tabula Muris Senis atlas (16 droplet-based and 12 Smart-seq2 fluorescence-activated cell sorting (FACS) sources). Cross-organ alignment of ageing programmes was quantified by genome-wide vector correlation, pre-ranked gene-set enrichment and a systemic ageing score.

Cochlear ageing showed weak but significant genome-wide concordance with systemic ageing (Spearman ρ 0.01–0.12), with lineage-matched pairs ranking among the highest. In contrast, systemic tissue ageing-up signatures, dominated by immune and inflammatory programmes, were depleted from the ageing-up rankings of cochlear non-sensory epithelial cells (normalised enrichment score −1.45, adjusted *p* = 0.032) and, for the immune-organ signatures, from cochlear macrophages. The cochlear macrophage ageing vector correlated positively with tissue-resident macrophages of peripheral organs (kidney ρ = 0.056, 95% confidence interval 0.016–0.063, *p* = 1.6×10^−13^; microglia ρ = 0.034, *p* = 1.4×10^−6^), negatively with bone-marrow and splenic macrophages, and not with lymphocytes. A systemic ageing score increased with age in spiral ganglion neurons, Schwann cells, inner hair cells, epithelial cells, and fibrocytes, but not in cochlear macrophages. These cell-type-resolved analyses support a compartmentalised view of cochlear ageing: non-immune compartments track systemic ageing weakly, the immune arm is reversed, and cochlear macrophages age along a tissue-resident rather than haematopoietic programme.

## Introduction

Ageing is a systemic process, and multi-organ single-cell atlases have begun to define its shared molecular programmes. The Tabula Muris Senis (TMS) atlas profiled more than 350,000 cells from 23 mouse organs across the lifespan and identified cell-type-specific manifestations of ageing hallmarks, including senescence, genomic instability, and altered immune function [1,2]. Complementary bulk RNA-sequencing of 17 organs showed that ageing gene-expression trajectories are broadly consistent across tissues, differing mainly in amplitude and onset, and that immune-cell activation is the most pronounced and earliest-detectable systemic ageing programme [3]. Together, these resources define a reference against which any individual organ’s ageing can now be measured.

The cochlea is absent from these atlases, although age-related hearing loss affects more than a quarter of adults aged over 65 years and ranks among the most prevalent sensory deficits of ageing [4,5]. A single-nucleus atlas of the ageing mouse inner ear recently described cell-type heterogeneity, hair-cell subtype-specific signatures, and macrophage activation in the aged cochlea [6]. Whether cochlear ageing programmes align with those of other organs, or whether the cochlea ages on its own terms, has not been examined. This question matters clinically: if cochlear ageing inherits systemic inflammatory programmes, anti-inflammaging strategies might be expected to extend to the inner ear; if not, organ-specific approaches are required.

Here we aligned the ageing cochlea against 28 systemic sources from the Tabula Muris Senis using three complementary and pre-registered statistics: genome-wide vector correlation without signature selection, pre-ranked gene-set enrichment with permutation testing, and a systemic ageing score built exclusively from systemic data. All analyses were accompanied by positive controls and platform sensitivity checks. We report evidence consistent with a compartmentalised cochlear ageing programme: non-immune compartments share a weak but significant trunk of systemic ageing, the immune/inflammatory arm is reversed, and cochlear macrophages age on a tissue-resident rather than haematopoietic programme.

## Methods

### Data sources

Mouse cochlear single-nucleus RNA-sequencing (snRNA-seq) data were obtained from the Gene Expression Omnibus (GEO) under accession GSE274279 [6], comprising nuclei from C57BL/6J mouse cochleae at 3, 12, and 24 months (mo; four 10x libraries; the two 24-mo libraries, 24M_Cochlea and 24M_Cochlea_BU, were reported as biological and technical replicates and were merged for the 24-mo group). Systemic reference data were obtained from the Tabula Muris Senis resource [1], comprising (i) droplet-based (10x) single-cell RNA-sequencing (scRNA-seq) for 16 tissues and (ii) Smart-seq2 FACS-enriched scRNA-seq for selected tissues (Thymus, Trachea, Pancreas, Skin, large intestine, Heart, brown (BAT), gonadal (GAT), mesenteric (MAT) and subcutaneous (SCAT) adipose tissue, Brain_Myeloid, Brain_Non-Myeloid). Both datasets are publicly available (see Data availability).

### Cochlear nucleus processing and clustering

Libraries were quality-controlled per sample (genes per nucleus 200–6,000; mitochondrial fraction <20%; total counts ≥500), yielding 13,988 nuclei. The original study reported 12,355 cochlear nuclei after the authors’ quality control [6]; the present thresholds retained 13,988 nuclei, and all downstream analyses used the independently quality-controlled set. After library-size normalisation to counts per 10,000 (CP10K) and log1p transformation, 3,000 highly variable genes were selected; principal component analysis (PCA; 30 components), a Gaussian-kernel neighbour graph, and Leiden clustering (resolution 0.5) [7] were computed with scanpy [8,9].

Clusters were annotated by canonical marker scores. Hair cells were identified by *Myo7a*/*Pou4f3*, with *Slc17a8*/*Kcnq4* marking inner hair cells (IHC) and *Ocm*/*Calb2* marking outer hair cells (OHC); spiral ganglion neurons (SGN) by *Tubb3*/*Map2*/*Rbfox3*; macrophages by *C1qa*/*C1qb*/*Cx3cr1*; Schwann cells by *Plp1*/*Mpz*; fibrocytes by *Col1a1*/*Dcn*/*Pdgfra*; and non-sensory epithelial cells by *Krt8*/*Krt18*/*Krt19*. These marker scores defined eight principal compartments (IHC, OHC, SGN, resident macrophages, Schwann cells, cochlear fibrocytes, non-sensory epithelial cells, and an *Apoe*-high mid-age activated state), plus a 24-mo–specific stressed-neuronal state analysed separately.

### Systemic tissue ageing signatures

For each TMS tissue, per-cell-type pseudobulk counts were computed by aggregating all nuclei of a cell type within each mouse (mouse-level biological replicates preserved). Pseudobulk counts were normalised to logCP10K, and ageing log-fold-change (LFC) per gene was computed as the difference of means between the old group (≥18 mo) and young group (≤3 mo), requiring ≥2 mice per group. Ageing signatures were defined per tissue × cell type by two pre-registered criteria: |LFC| ≥ 0.5 and directional consistency ≥ 0.80 across all old–young mouse pairs. We pre-registered this effect-size/consistency criterion instead of FDR-controlled differential expression because 2–3 mice per age group preclude FDR-level inference; signatures are used for scoring and enrichment, not for mechanistic claims.

The cochlear data do not contain mouse-level replicates (one merged library per age). Cochlear ageing LFC vectors were therefore estimated at the single-cell level with bootstrap resampling (B = 100–200 stratified by library) and reported with bootstrap z-scores (|z| ≥ 2 with detection ≥ 3–5% used for signature membership); no parametric significance tests were applied at this step (declared limitation).

### Cross-tissue alignment statistics

Three complementary, pre-registered statistics were computed. First, genome-wide vector concordance: Spearman correlation between cochlear cell-type ageing LFC vectors and systemic tissue/cell-type ageing LFC vectors over all shared genes (*n* = 17,588–19,610). Second, gene-set enrichment analysis (GSEA): pre-ranked (weighted Kolmogorov–Smirnov running sum, |LFC| weighting, 1,000 gene-set permutations; permutation nulls precomputed per ranked-list size; Benjamini–Hochberg correction within each cochlear cell type [10]). Both ageing-up and ageing-down signatures were tested (8 cochlear compartments × 16 systemic sources × 2 directions = 256 comparisons). Third, systemic ageing score trajectories. A consensus systemic signature (the top 150 up- and top 150 down-regulated genes by mean ageing LFC across the 10 droplet tissues;

TMS-only information, avoiding cochlear circularity) was scored per cochlear nucleus as the mean logCP10K of systemic-up genes minus that of systemic-down genes. Trajectories across 3, 12, and 24 mo were summarised per cell type (median) with cell-level Spearman trend tests.

Control analyses, all pre-registered before interpreting the cochlea–system comparisons, were: (i) TMS–TMS positive control (tissue signatures projected onto other tissues’ rankings); (ii) detection-rate bias check for anti-enriched gene sets; (iii) median-centering of cochlear rankings to exclude global-shift artifacts; (iv) platform sensitivity analysis (droplet vs Smart-seq2 FACS systemic sources).

### Composition, noise, and pathway analyses

Cell-type composition across ages was compared with chi-square tests on cell-type × age contingency tables and two-proportion z-tests (3-mo vs 24-mo). Expression noise was quantified as the median cell-to-cell coefficient of variation over genes detected in ≥5% of nuclei within each compartment × age. Splicing-factor changes were screened across 36 canonical splicing-factor genes (*Rbm25*, Srsf, Hnrnp, U2af, Tra2, Rbfox, Ptbp, Qk, Nova, Mbnl, Celf, Khdrbs, Elavl families) using the bootstrap z criterion. Gene-set enrichment of reversed gene sets (systemic ageing-up genes that decline in a cochlear compartment) used clusterProfiler [11] with org.Mm.eg.db for Gene Ontology (GO) Biological Process enrichment, BH-adjusted; KEGG online queries were unavailable and were replaced by GO-only enrichment. Cell-type composition and expression noise are reported in Results.

### Statistics

All decision criteria were pre-registered: |LFC| ≥ 0.5 with ≥ 0.80 directional consistency for systemic signatures, |bootstrap z| ≥ 2 with ≥ 3–5% detection for cochlear signatures, and ≥ 15 mappable genes for enrichment. Because the cochlea is represented by one merged library per age, cochlear-level uncertainty was quantified by stratified bootstrap resampling (percentile 95% confidence intervals, B = 100–200) rather than parametric or mouse-level inference; mouse-level biological replication is preserved for all TMS-derived estimates. As a built-in replication check on the cochlear single-merged-library design, the two 24-mo libraries were additionally analysed separately (see Results). Enrichment significance was assessed by permutation (1,000 gene-set permutations with precomputed nulls) under Benjamini–Hochberg control within each cochlear compartment; composition contrasts used chi-square tests and two-proportion z-tests, and trajectory trends used cell-level Spearman correlation. Dispersion is reported as median cell-to-cell coefficients of variation (expression noise) and percentile bootstrap confidence intervals for correlation coefficients.

### Software and reproducibility

Python 3.13 with scanpy [8], anndata/scverse [9], numpy, pandas, scipy [12], statsmodels, and matplotlib [13] was used. All analysis scripts, intermediate tables, and permutation settings are publicly archived at Zenodo (https://doi.org/10.5281/zenodo.24217326); every figure regenerates from the archived tables.

### Use of large language models

The manuscript text was drafted and revised with LLM assistance under the authors’ direction; all claims, statistics, and references were verified against primary sources by the authors, who take full responsibility for the integrity and accuracy of the work. LLMs were not used to generate data, images, or scientific conclusions.

## Results

### A single-nucleus atlas of the ageing mouse cochlea resolves eight principal compartments

Quality control retained 13,988 cochlear nuclei (3 mo, *n* = 2,868; 12 mo, *n* = 5,467; 24 mo, *n* = 5,653 after merging 24M libraries). Leiden clustering resolved 17 clusters, annotated into eight principal compartments. An *Apoe*-high activated state peaked at 12 mo, independently reproducing the reported biphasic *Apoe* trajectory [6]. A 24-mo–specific stressed-neuronal state was analysed separately; its top genes (*Gm42418, Gabra2, Banf2*) matched the leading genes of the whole-cochlea ageing-up signature.

### Cochlear ageing preserves a weak but significant genome-wide concordance with systemic ageing

Genome-wide Spearman correlation between cochlear cell-type ageing log-fold-change (LFC) vectors and systemic ageing vectors was positive in the large majority of pairs (ρ 0.01–0.12; strongest pairs *p* < 10^−20^, *n* = 17,588–19,610 shared genes; Fig. 1). Lineage-matched pairs ranked among the highest in their rows (epithelial × trachea 0.094; Schwann × pancreas 0.115 and trachea 0.108; inner hair cells (IHC) × Brain_Non-Myeloid 0.077), indicating that the concordance reflects biology rather than platform artifacts.

**Fig. 1.**
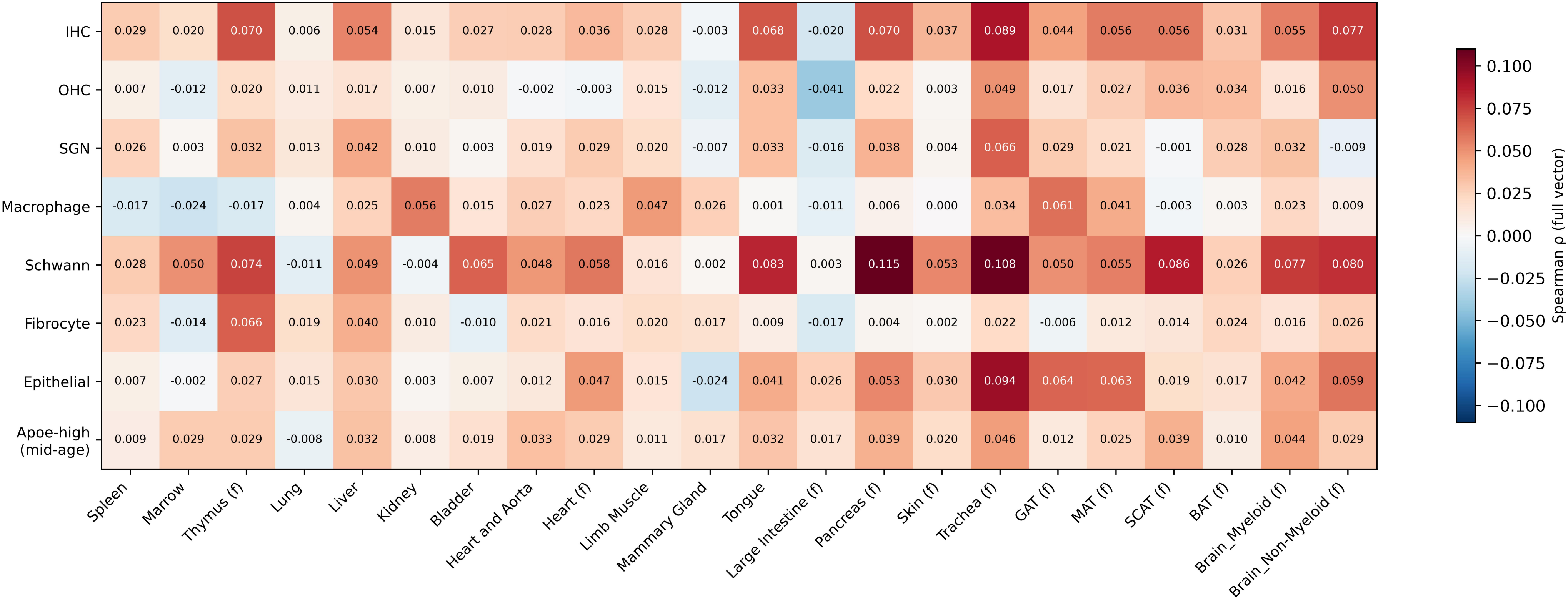
Genome-wide ageing concordance between cochlear cell types and systemic tissues. Spearman correlation of ageing log-fold-change vectors (24-mo vs 3-mo) over all shared genes (*n* = 17,588–19,610), computed without signature selection, between each cochlear cell type (rows) and each systemic source (columns; 10 droplet-based tissues, unlabelled, and 12 Smart-seq2 FACS sources, labelled (f); aggregated across cell types per mouse). The colour scale is symmetric around zero (white). For the macrophage row, droplet-tissue columns show macrophage-specific systemic vectors (or leucocyte-specific where no macrophage lineage is annotated), and the Brain_Myeloid column shows the whole-myeloid tissue vector. Lineage-matched pairs rank among the highest within rows (e.g., epithelial × trachea 0.094; Schwann × pancreas 0.115; IHC × Brain_Non-Myeloid 0.077).

### The immune/inflammatory arm of systemic ageing is reversed in the ageing cochlea

In contrast to the weak genome-wide concordance, systemic tissue ageing-up signatures were depleted from the top of cochlear ageing rankings. Across all 256 comparisons, the only Benjamini–Hochberg (BH)-significant result was an anti-enrichment: the spleen ageing-up signature was depleted from the ageing-up ranking of cochlear non-sensory epithelial cells (normalised enrichment score (NES) −1.45, permutation *p* = 0.001, BH = 0.032; Fig. 2). Nominal anti-enrichments were also observed for spleen/liver signatures in cochlear macrophages (*p* = 0.015–0.025). Median-centering, detection-bias checks, and the TMS–TMS positive control (strong enrichment, NES 1.1–2.3, with most BH-adjusted *p* < 0.05) excluded methodological and detection artifacts.

**Fig. 2.**
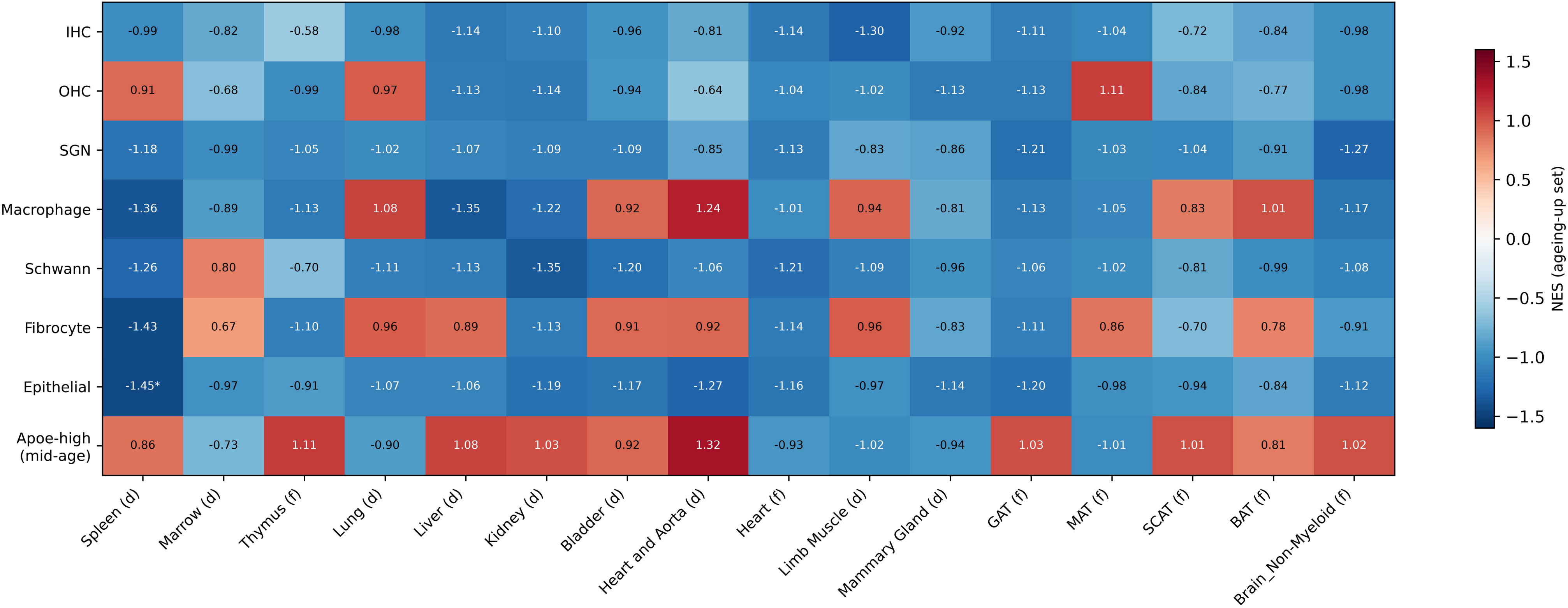
Cross-organ alignment of systemic ageing signatures onto cochlear cell-type ageing rankings. Normalised enrichment scores (NES) of pre-ranked GSEA in which the ageing-up signature of each systemic source (columns; suffixes (d)/(f) denote droplet-based or Smart-seq2 FACS platform) was projected onto the ageing ranking (24-mo vs 3-mo log-fold change) of each cochlear cell type (rows; IHC, inner hair cells; OHC, outer hair cells; SGN, spiral ganglion neurons). Shown are the 16 sources with at least 15 genes mappable to the cochlear gene space; the remaining six sources were omitted (tongue (d) and Brain_Myeloid (f) signatures had fewer than 15 mappable genes and were null in genome-wide vector analysis; Pancreas, Skin, Trachea, and large intestine are represented in TMS by both platforms; their FACS versions were reserved for the platform-sensitivity analysis and excluded from the main matrix to avoid platform-confounded duplication). Positive NES indicates concordance; negative NES indicates anti-enrichment. Asterisks mark Benjamini–Hochberg-adjusted permutation *p* < 0.05 within each cochlear compartment (256 comparisons over these 16 sources). The single significant comparison (epithelial × spleen, NES −1.45, BH = 0.032) is anti-enrichment.

#### Platform sensitivity

All primary analyses were repeated with the 12 FACS sources included (18 systemic sources designed; 256 comparisons for the 16 whose signatures contained at least 15 mappable genes). The epithelial × spleen anti-enrichment remained the only Benjamini– Hochberg-significant comparison (NES −1.447, BH = 0.032), and the epithelial row was negative across all 16 mappable sources (NES −0.85 to −1.45). Genome-wide concordance values were reproduced with FACS-derived systemic vectors (Schwann × pancreas 0.115; epithelial × trachea 0.094; IHC × Brain_Non-Myeloid 0.077). One exception is noted in Discussion: fibrocyte vectors were null against all FACS sources while positive against droplet lung.

#### Library-level replication

As a built-in check on the single-merged-library design, the two 24-mo libraries were analysed separately. Their genome-wide ageing log-fold-change vectors correlated at ρ = 0.94 with each other and at ρ ≥ 0.96 with the merged vector (*n* = 9,707 genes), the direction of 3-mo-to-24-mo change in the systemic ageing score was concordant across the two libraries in all 15 evaluable Leiden clusters (two clusters lacked 3-mo nuclei and were not testable), and the macrophage trajectory (12-mo peak, 24-mo decline) was reproduced within each library (cell-level ρ = −0.04 and −0.07).

### Composition, expression noise, and splicing-factor changes across cochlear compartments

Cell-type composition shifted strongly with age across the nine composition categories retained after excluding the 24-mo–specific state (the seven principal compartments, the *Apoe*-high state, and a residual Other class; global chi-square = 1,633.7, degrees of freedom (df) = 16, *p* < 10^−300^; Fig. 3a). The *Apoe*-high state was biphasic at the composition level (2.34%, 14.69%, and 0.80% of all nuclei at 3, 12, and 24 mo; 24-mo vs 3-mo *z* = −5.90), quantifying at the composition level the biphasic trajectory noted above [6]. A 24-mo–specific stressed-neuronal state expanded from 0.45% to 56.25% of all nuclei. Because each age is represented by a single merged library, nucleus-capture efficiency and library-level composition effects cannot be separated from biological expansion for this compartment; we therefore report the shift descriptively rather than as a magnitude of proliferation or activation. The same caveat applies in weaker form to all composition estimates in this section. The fractions of Schwann cells (16.35% → 0.57%), outer hair cells (OHC; 5.20% → 0.41%), inner hair cells (5.30% → 1.19%), and fibrocytes (50.07% → 23.81%) declined, macrophages declined modestly (2.27% → 1.61%, *z* = −2.14, *p* = 0.033), and the epithelial fraction was stable (*p* = 0.14).

**Fig. 3.**
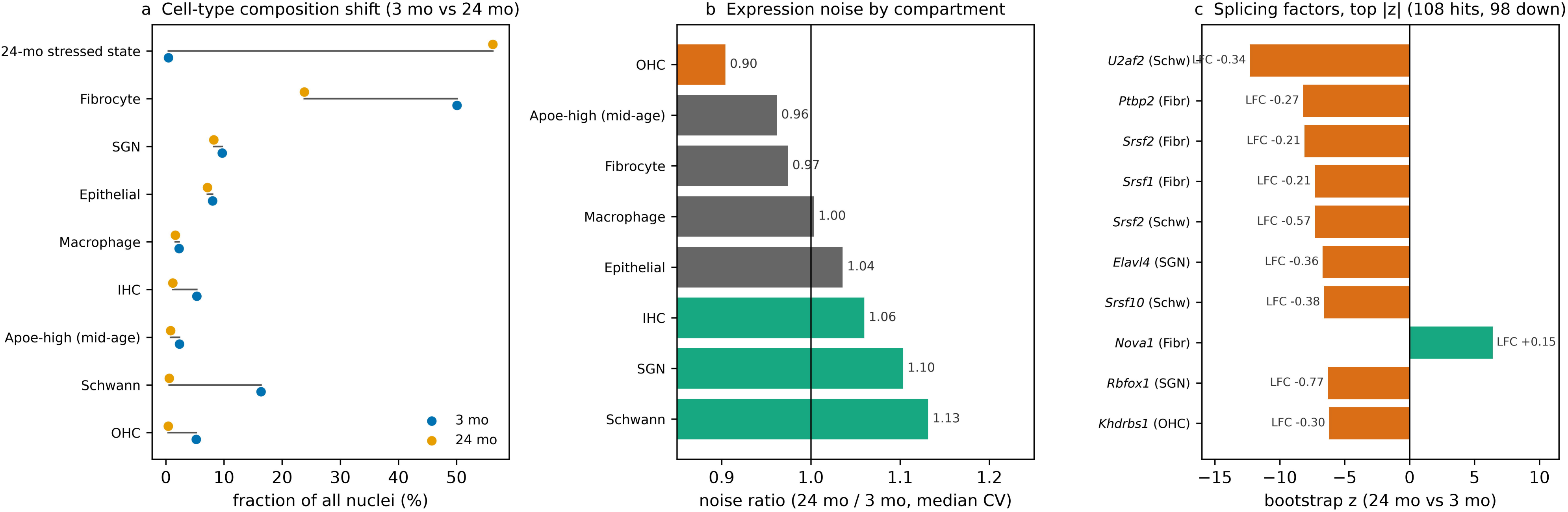
Composition, expression noise, and splicing-factor changes. (a) Cell-type composition of all cochlear nuclei at 3 mo (blue) and 24 mo (orange); global chi-square = 1,633.7, *df* = 16, *p* < 10^−300^ across the nine composition categories retained after excluding the 24-mo–specific state (seven principal compartments, *Apoe*-high state, and a residual Other class). (b) Expression-noise ratio (median cell-to-cell coefficient of variation, 24-mo / 3-mo) per compartment; teal = increase, vermillion = decrease, grey = stable. (c) Splicing-factor changes with |bootstrap z| ≥ 6, by compartment; vermillion = down-regulation, teal = up-regulation. In total, 108 splicing-factor changes (|z| ≥ 2) were detected, of which 98 were down-regulations.

Per-cell-type expression noise (median cell-to-cell coefficient of variation across detected genes) rose with age in spiral ganglion neurons (SGN; 24-mo/3-mo ratio 1.10), Schwann cells (1.13), IHC (1.06), and epithelial cells (1.04), was flat in macrophages (1.00), fibrocytes (0.97), and the *Apoe*-high state (0.96), and decreased in outer hair cells (0.90), consistent with selective loss of an OHC subpopulation leaving a more homogeneous surviving population (Fig. 3b).

A screen of 36 splicing-factor genes identified 108 age-associated changes (|bootstrap z| ≥ 2) across all eight compartments, of which 98 were down-regulations (Fig. 3c). Representative examples include *U2af2* in Schwann cells (LFC −0.34, *z* = −12.3), *Rbfox1* in SGN (−0.77, *z* = −6.3), *Elavl4* in SGN (−0.36, *z* = −6.7), and *Khdrbs1* in OHC (−0.30, *z* = −6.2). In contrast to the broad decline of the general splicing machinery, *Rbm25* itself was specifically up-regulated in IHC (+0.42, z = +2.4), epithelial cells (+0.55, z = +5.1), and fibrocytes (+0.22, z = +4.7). This pattern connects the cochlear ageing alignment to the mechanism reported for inflammatory hair-cell degeneration, in which suppressing *Rbm25* rescued degenerating hair cells [6].

### Cochlear resident macrophage ageing aligns with a tissue-resident, not haematopoietic, programme

The cochlear macrophage ageing vector correlated positively with tissue-resident macrophages of peripheral organs (kidney ρ = 0.056, 95% confidence interval (CI) 0.016–0.063, *p* = 1.6×10^−13^; limb muscle 0.047, 95% CI 0.000 to 0.072; mammary gland 0.026, 95% CI −0.003 to 0.031), negatively with haematopoietic-centre macrophages (bone marrow −0.024, 95% CI −0.038 to 0.008, *p* = 1.5×10^−3^; spleen −0.017, 95% CI −0.039 to 0.012, *p* = 0.024), and showed no correlation with lymphocytes (12 lineages, |ρ| ≤ 0.022). Only the kidney correlation had a bootstrap confidence interval that excluded zero; the remaining directions were consistent but their confidence intervals included or touched zero. The microglial ageing vector (Brain_Myeloid, 13,130 microglia) correlated positively (ρ = 0.034, *p* = 1.4×10^−6^) but fell below the strongest resident-macrophage alignment (kidney, ρ = 0.056); we therefore report this pattern without ranking the strengths of these alignments.

### The systemic ageing clock ticks in non-immune cochlear cells but stalls in macrophages

Scoring cochlear nuclei with the TMS-derived consensus systemic signature showed significant age-increasing trajectories in SGN (ρ = 0.268, *p* = 3.9×10^−23^), Schwann cells (0.223), IHC (0.215), epithelial cells (0.165), and fibrocytes (0.076) (Fig. 4). Cochlear macrophages were the only compartment with a non-positive trajectory (ρ = −0.044, ns); their median score peaked at 12 mo and declined by 24 mo. OHC and the *Apoe*-high state were also non-significant but directionally positive (0.045 and 0.004), the latter consistent with its biphasic definition.

**Fig. 4.**
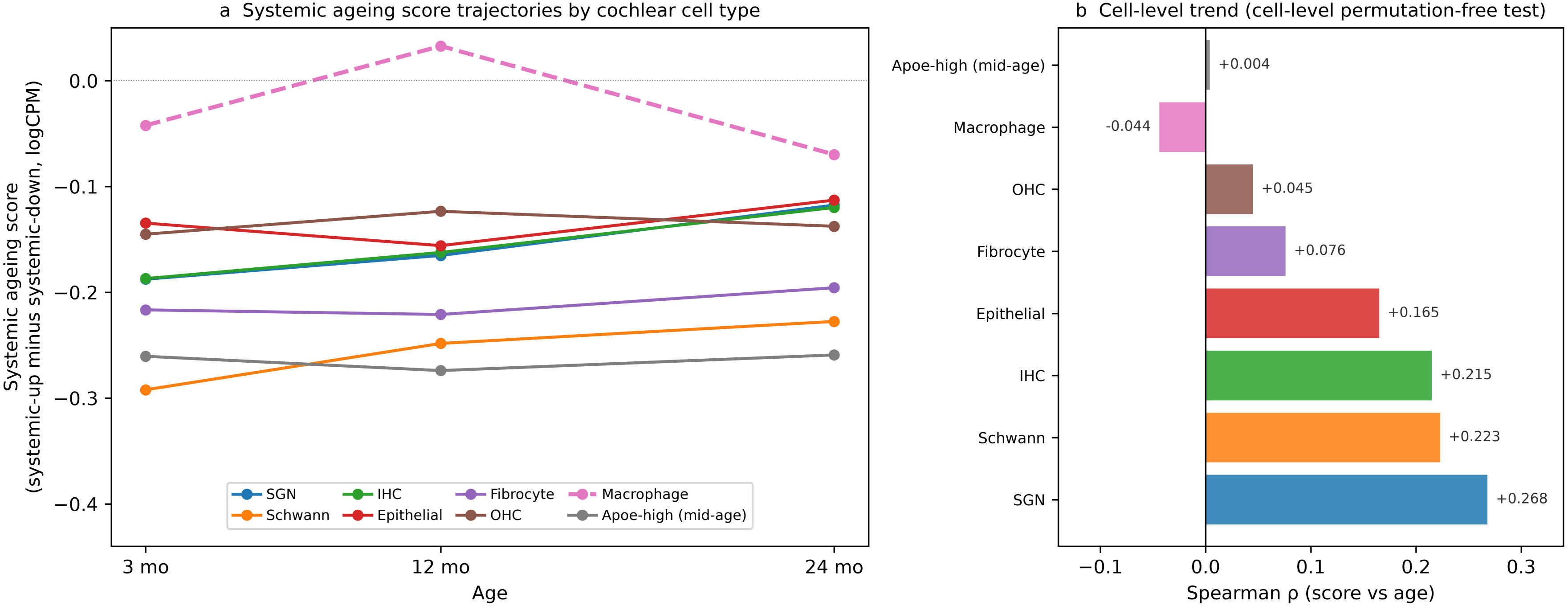
Systemic ageing score trajectories across cochlear compartments. (a) Median systemic ageing score (mean logCP10K of the consensus systemic ageing-up signature minus the systemic ageing-down signature, both derived from TMS tissues only) at 3, 12, and 24 mo for each cochlear cell type. (b) Cell-level Spearman correlation between the systemic ageing score and age for each compartment (cell-level trend test). Correlations were significant for SGN, Schwann cells, IHC, epithelial cells, and fibrocytes (each *p* < 0.001; fibrocytes *p* = 2.6 × 10^−8^) but not for OHC, macrophages, or the *Apoe*-high state.

## Discussion

We aligned a single-nucleus atlas of the ageing mouse cochlea against the Tabula Muris Senis multi-organ atlas and found evidence consistent with a compartmentalised cochlear ageing programme. Across the genome, ageing transcriptional changes in cochlear cells correlated weakly but significantly with systemic tissue ageing, and the lineage-matched pairs ranked among the highest within each cochlear compartment. At the same time, the systemic ageing-up signatures were depleted from the ageing rankings of cochlear epithelial cells; in cochlear macrophages, the immune-organ signatures were depleted (NES −0.89 to −1.36) while adipose-, vessel-, and muscle-derived signatures were modestly enriched (+0.83 to +1.24), consistent with shared resident-macrophage programmes. These gene sets define tissue ageing in the atlas and are dominated by immune and inflammatory programmes. The macrophage compartment was the sharpest example: its ageing programme correlated with tissue-resident macrophages of peripheral organs and with microglia, diverged from bone-marrow and splenic macrophage ageing, and showed no relationship to lymphocyte ageing. A systemic ageing score built exclusively from systemic data increased with age in spiral ganglion neurons, Schwann cells, inner hair cells, epithelial cells, and fibrocytes, but not in cochlear macrophages. These results indicate that the cochlea neither ages as an isolated organ nor simply inherits systemic ageing: it shares the metabolic and structural core of systemic ageing while keeping its immune arm compartmentalised.

The reversed immune arm is notable because both the systemic and the cochlear data independently describe immune activation with age. Across TMS tissues, immune-cell activation is the most pronounced and earliest-detectable ageing programme, first emerging in white adipose depots at middle age [3]. In the aged inner ear, macrophage activation and inflammatory niches are well documented [6]. Our results do not contradict these observations but separate two readily conflated phenomena. Local macrophage activation within the aged cochlea occurs, but the direction of gene-expression change that defines ageing in spleen and liver, largely haematopoietic-centre and monocyte-lineage programmes, does not carry into the cochlea.

Cochlear macrophages instead share their ageing programme with tissue-resident macrophages of peripheral organs. This pattern is consistent with the developmental biology of the resident macrophage lineage: most tissue macrophages derive from embryonic precursors, are seeded before birth, and maintain themselves in adults through local self-renewal rather than replacement by blood monocytes [14], and the cochlea hosts such resident populations in the spiral ganglion, spiral ligament, and stria vascularis [15,16]. The ageing cochlea thus mounts local inflammation, but its macrophages do not adopt the haematopoietic ageing programme that dominates systemic immune ageing. Pathway dissection of the reversed gene sets localises this reversal to concrete machinery: the systemic ageing-up genes that decline in the ageing cochlear epithelium are enriched for leukocyte aggregation and leukocyte cell–cell adhesion programmes (BH-adjusted *p* = 4.1×10^−4^), and those that decline in cochlear macrophages are enriched for leukocyte migration and mitotic nuclear-division programmes (BH-adjusted *p* = 1.1×10^−4^ and 8.0×10^−5^, respectively). The aged cochlea thus disengages the adhesion and migration machinery through which systemic immune cells would be recruited. Its macrophages suppress the proliferative programmes that define haematopoietic ageing. In a parallel screen, the general splicing machinery declined across compartments while the *Rbm25* branch rose specifically in sensory and epithelial cells. This connects the cochlear alignment to the splicing-based mechanism reported for inflammatory hair-cell degeneration [6].

Our findings also connect to the recently described “gut–ear axis”. Faecal microbiota transplantation from patients with active ulcerative colitis disrupts the blood-labyrinth barrier (BLB), activates cochlear MyD88/NF-κB signalling, recruits IBA-1-positive macrophages, and impairs auditory function, whereas microbiota from remission-phase patients is relatively protective [17]. A recent review proposed metabolic, inflammatory, and neural routes for this axis and noted that its contribution to presbycusis remains unvalidated [18]. Our results add the dimension that this literature lacks. The inflammatory signals that cross the BLB in experimental colitis are pathological, acute perturbations; the physiological ageing programmes of systemic immune organs, by contrast, appear to be filtered out of the ageing cochlea (Figs. 1 and 2). We therefore propose that the BLB acts as a selective gatekeeper: pathological immune signals penetrate and engage cochlear macrophages through MyD88/NF-κB, whereas physiological haematopoietic ageing programmes do not. Testing this hypothesis directly, by transplanting microbiota from aged into young mice and profiling cochlear nuclei, would establish whether systemic ageing signals penetrate the BLB and link the present findings to the causal toolkit of the gut–ear literature.

The weak genome-wide concordance warrants comment on effect size. The ρ values we report (0.01–0.12) are small, as might be expected for cross-organ ageing comparisons at cell-type and platform resolution. Every structural feature of the matrix supported a biological interpretation: lineage-matched pairs ranked among the highest, the epithelial anti-enrichment was uniform across all 16 mappable systemic sources, and TMS–TMS positive controls showed that the enrichment framework detects strong concordance when it exists. One discrepancy requires explicit mention: fibrocyte ageing vectors were weakly positive against droplet-derived systemic sources but null against all FACS sources. This platform contrast was not observed for any other cochlear compartment and most likely reflects the combination of full-transcript Smart-seq2 coverage with the fibrocyte extracellular-matrix programme, although a modality-specific contribution cannot be excluded. A second unexplained pattern was the absence of concordance between spiral ganglion neurons and brain non-myeloid cells despite shared neuronal identity; the SGN snRNA detection profile may underlie this exception.

Four limitations frame the interpretation. First, the cochlear data are single-nucleus RNA-sequencing (snRNA-seq) data while the systemic references are predominantly single-cell RNA-sequencing (scRNA-seq) data; nuclear retention biases cytoplasmic and secreted transcripts. Our detection-rate checks found no under-detection of the anti-enriched gene sets, and genome-wide positive concordance argues against a globally suppressive artifact, but the modality difference cannot be eliminated and should be borne in mind in any mechanistic interpretation. Second, the cochlear data comprise one merged library per age; we therefore used cell-level bootstrap resampling and relied on pre-declared non-parametric criteria rather than mouse-level statistical inference. Third, the C57BL/6J background carries early-onset progressive hearing loss, and the 24-mo outer and inner hair cell compartments contained few nuclei; the hair-cell-specific results should be read as exploratory. Finally, all findings are associative; the alignment identifies programmes, not mechanisms.

These findings revise the default assumptions about how the cochlea ages. Before multi-organ atlases, the cochlea was studied largely on its own terms, as an isolated sensory organ; once the atlases appeared, the implicit expectation became that every organ follows the systemic ageing programmes they define. Our data support both positions in part: the non-immune compartments of the cochlea do track systemic ageing weakly but significantly, while the immune arm runs counter to it. The compartmental pattern we describe (shared trunk, reversed immune arm, resident-lineage macrophage ageing) is therefore not a choice between the two earlier positions but a structural statement about where the cochlea sits on the systemic ageing map.

At the clinical level, the implication concerns which interventions can be expected to reach the inner ear. Systemic strategies marketed toward inflammaging [19] (anti-inflammatory diets, systemic anti-inflammatory agents, NAD+-targeting supplements) cannot be assumed to protect hearing through simple immune-programme sharing, because the cochlear immune compartment does not follow the systemic inflammatory ageing programme. Conversely, systemic health management targeting the metabolic and vascular components of ageing remains meaningful for the trunk of cochlear ageing. If the cochlear macrophage ages on a tissue-resident programme, future strategies to delay cochlear ageing should target resident-lineage pathways specifically rather than importing anti-inflammatory logic from systemic immunology. Beyond the inner ear, age-related hearing loss ranks among the largest potentially modifiable contributors to dementia risk [20]. Our data add a compartmentalised caveat to this association: if the cochlea tracks systemic metabolic and structural ageing while filtering haematopoietic inflammatory ageing, then the hearing-loss–dementia link is unlikely to be modified by systemic anti-inflammatory strategies alone, and audiometric decline may merit evaluation as an accessible peripheral readout of compartmentalised ageing.

Methodologically, the alignment pipeline itself is a transferable contribution. Multi-organ atlases do not cover every organ; neither the Tabula Muris Senis nor human atlas efforts include the inner ear. The three-statistics alignment used here (genome-wide vector correlation, pre-ranked enrichment with permutation testing, and a systemic-only scoring trajectory), together with positive controls, bias checks, and platform sensitivity analysis, can be applied to any organ absent from existing atlases, in mouse or human. Extending the same comparison to long-lived human cohorts with preserved hearing, such as centenarian studies, would test whether strong auditory preservation corresponds to an even more pronounced macrophage compartmentalisation.

## Acknowledgments

The authors thank the Tabula Muris Senis consortium and the authors of GSE274279 for making the data publicly available.

## Statements and Declarations

### Funding

This research received no specific grant from any funding agency in the public, commercial, or not-for-profit sectors.

### Competing Interests

The authors declare no conflicts of interest.

### Ethics

No human participants or animals were directly used in this study; all data are from previously published, ethically approved public resources (GSE274279; Tabula Muris Senis).

### Author contributions

Zhengyi Tang: Conceptualization, Methodology, Software, Formal analysis, Data curation, Writing – original draft, Writing – review & editing, Visualization. Ting Liao: Formal analysis, Investigation, Data curation, Writing – review & editing. Qiuping Lin: Validation, Investigation, Data curation, Writing – review & editing. Zhengyi Tang and Ting Liao contributed equally to this work.

### Data availability

Cochlear snRNA-seq data are available at GEO under accession GSE274279 [6]. Tabula Muris Senis data are available at figshare (https://doi.org/10.6084/m9.figshare.8273102.v2) and GEO (GSE132042) [1]. All analysis scripts and intermediate tables are publicly archived at Zenodo (https://doi.org/10.5281/zenodo.24217326).

## Notes

https://www.ncbi.nlm.nih.gov/geo/query/acc.cgi?acc=GSE274279

https://doi.org/10.6084/m9.figshare.8273102.v2

https://doi.org/10.5281/zenodo.24217326

### Competing Interest Statement

The authors have declared no competing interest.

